# HiFi-ST: High-Fidelity Reconstruction of Continuous Spatial Transcriptomic Expression Fields via Conditional Neural Fields

**DOI:** 10.64898/2026.06.29.735170

**Authors:** Lei Tang, Wenshuai Han, Xiao Yang, Xiaozhou Chen, Huamei Li

**Author notes:** Correspondence: Huamei Li, Xiaozhou Chen.

## Abstract

Spatial transcriptomics characterizes tissue-scale gene expression patterns, yet its observations are sparse discrete samples of an underlying continuous molecular field, leading to spatial aliasing and sub-resolution information loss. Existing methods usually formulate this task as spot-level point regression, making it difficult to capture both expression continuity and the regional nature of observation. Here, we propose HiFi-ST, a conditional neural field framework for continuous spatial transcriptomics modeling. HiFi-ST formulates spatial gene expression prediction as continuous expression field learning, models each spot as a regional observation over a finite support domain, approximates local integration through Monte Carlo sampling, and integrates multiscale tissue feature extraction with FiLM-based conditional modulation to improve modeling of complex spatial heterogeneity and consistency with the underlying measurement process. Systematic evaluation on three independent datasets (HER2+, cSCC, and Alex_NatGen) showed that HiFi-ST outperformed mclSTExp, BLEEP, THItoGene, His2ST, and HisToGene on key metrics. On HER2+, HiFi-ST achieved an average PCC improvement of 65.1% and an average MSE reduction of 40.9%; on cSCC, PCC improved by 10.2% and MSE decreased by 51.2%; on Alex_NatGen, PCC improved by 80.0% and MSE decreased by 16.3%. In addition, the learned multiscale tissue representations supported downstream spatial immunoanalysis, including assisted identification of candidate TLS regions. Overall, HiFi-ST provides a unified framework bridging discrete measurements and continuous expression field reconstruction for tumor microenvironment analysis and spatial immune structure characterization.

## Introduction

Gene expression in biological tissues is intrinsically continuous, whereas current spatial transcriptomics technologies discretely sample the underlying molecular field because of physical capture-array spacing, causing spatial aliasing and sub-resolution information loss ^[1–3]^. Predicting spatial gene expression from H&E images has therefore become an important strategy for mitigating these limitations ^[4–6]^. Representative methods include Vision Transformer-based sequence modeling, graph neural network-based topological modeling, and multimodal contrastive learning approaches such as HisToGene ^[7]^, His2ST ^[8]^, BLEEP ^[9]^, and mclSTExp ^[10]^. However, these methods still follow a discrete spot-level supervision paradigm and do not explicitly model expression continuity or regional observation ^[11, 12]^.

To address this issue, we propose HiFi-ST, a conditional neural field ^[13]^ framework for continuous spatial transcriptomics modeling. HiFi-ST reformulates spatial gene expression prediction as continuous expression field learning, models each spot as a regional observation over a finite support domain ^[14]^, and approximates local integration through Monte Carlo sampling. It further combines multiscale tissue feature extraction with FiLM-based conditional modulation ^[15]^ to improve modeling of complex spatial heterogeneity.

Our contributions are threefold: (i) reformulating spatial gene expression prediction from discrete point-wise regression to continuous expression field learning; (ii) explicitly introducing regional observation modeling to better match the measurement mechanism; and (iii) integrating multiscale conditional modulation with conditional neural field decoding for high-fidelity reconstruction of continuous expression patterns in complex tissues.

## Materials and Methods

### HiFi-ST Framework

Each tissue section is associated with an H&E image *I* and a set of sequencing locations *x_i_* with known spatial positions. HiFi-ST takes multiscale histological images and spatial coordinates as input, reconstructs gene expression in continuous space, and obtains spot-level predictions through regional sampling and aggregation. The framework consists of four steps: data preprocessing, multiscale feature extraction, conditional modulation, and random-sampling-based prediction and aggregation.

### Data preprocessing

Given N spatial locations and G genes, the raw count matrix is denoted as:

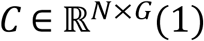

where *C_ij_* represents the raw transcript count of the j-th gene at the i-th spatial location.

Highly variable gene (HVG) selection was first performed to reduce low-information noise and improve cross-sample consistency. Within each tissue section, library-size normalization and log transformation were applied, followed by HVG identification using Scanpy; stable candidate genes were then integrated across sections ^[10]^. We used 1000 HVGs as the default upper limit, although the final number of genes included for prediction could be smaller because of cross-section gene overlap and data-quality constraints (Supplementary Table S1).

After defining the HVG subspace, CPM-like normalization and log transformation were applied to obtain the expression matrix:

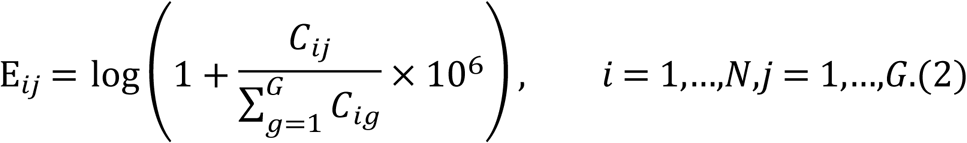

Where 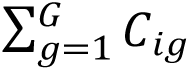denotes the total count of the i-th spot. The resulting normalized expression matrix is given by:

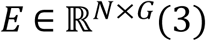

This matrix is used as the regression supervision signal for the model.

The spatial coordinate *x_i_* ∈ ℝ^2^is defined in the pixel coordinate system of the tissue section and is used to extract local image patches aligned with the corresponding spot from the H&E image. To enhance the capacity for continuous spatial modeling, the model further incorporates spatial coordinate embeddings based on Fourier positional encoding. In addition, to model the global expression structure, we used offline precomputed PCA results to reduce the dimensionality of the normalized expression matrix and retained the first 8 principal components as low-dimensional expression embeddings:

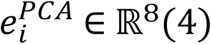

This embedding serves as an auxiliary conditional prior. The shared global condition vector is denoted as:

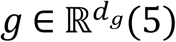

The optional expression PCA branch first projects *e_i_^PCA^* into the conditional space, with dropout applied during the mapping process. The projected representation is then fused with the shared global condition vector to form the final conditional vector:

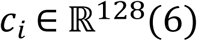

where *c_i_* denotes the conditional vector used in subsequent conditional feature modulation and conditional neural field prediction.

### Multi-scale feature extraction

To capture tissue morphological information at different spatial scales, HiFi-ST constructs three input fields of view around each spatial location *x_i_*: 112×112, 224×224, and 448×448.

Specifically, the 224×224 image patch is derived from a pre-cropped patch aligned with the spot coordinates; the 112×112 image patch is obtained by downsampling the 224×224 patch; and the 448×448 image patch is dynamically cropped from the full-resolution H&E section centered at *x_i_*. Therefore, only the 448×448 scale relies on dynamic cropping from the whole tissue section, whereas the other scales are obtained from patch-level data.

Each image patch *I_s_*_,*i*_ at scale *s* is fed into its corresponding convolutional neural network encoder Φ*_s_* to obtain the visual feature representation:

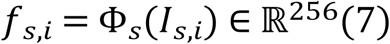

where Φ*_s_* denotes the image encoder corresponding to scale *s*, and *f_s_*_,*i*_ denotes the visual feature at that scale.

After obtaining the features at each scale, the model first performs conditional modulation, and then conducts multiscale fusion using learnable scale weights α*_s_*:

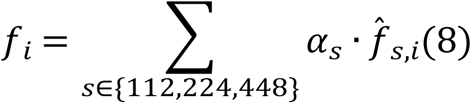

Where 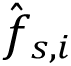 denotes the conditionally modulated feature, and α*_s_* is a learnable parameter initialized with a uniform value and jointly optimized during training. In this study, no softmax normalization constraint is imposed on α*_s_*; therefore, the sum of the weights across scales is not required to be strictly equal to 1.

### Conditional feature modulation

HiFi-ST adopts Feature-wise Linear Modulation (FiLM) to inject contextual priors into visual features. For each scale *s*, the conditional vector *c_i_* is mapped to modulation parameters:

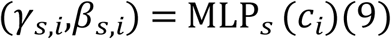

which are then applied channel-wise to the feature *f_s_*_,*i*_:

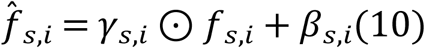

where *γ_s_*_,*i*_ and *β_s_*_,*i*_ denote the scaling and shifting parameters, respectively, and 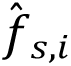 denotes the scale-specific feature after conditional modulation. The modulated multiscale features are then fed into the multiscale fusion module for aggregation.

### Prediction and Aggregation Based on Random Sampling

To improve robustness to spatial noise and minor registration errors, HiFi-ST performs random multi-point sampling within the neighborhood of each spatial location and approximates the expected regional observation by mean aggregation. For each location *x_i_*, M perturbations are sampled from an isotropic Gaussian distribution:

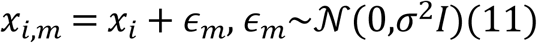

where σ = 0.1 and M=16. For each sampled coordinate *x_i_*_,*m*_, its spatial representation is constructed as:

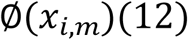

where ∅(·) denotes the positional encoding function. When Fourier positional encoding is enabled, ∅ corresponds to the Fourier feature mapping; otherwise, the raw coordinates are used directly.

Subsequently, the fused multiscale visual feature *f_i_* is concatenated with the spatial representation to form the neural field input. If the spatial PCA branch is enabled, the precomputed spatial PCA feature *u_i_* is further incorporated:

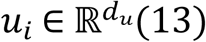

where *d_u_* denotes the dimensionality of the spatial PCA feature. The neural field input is then represented as:

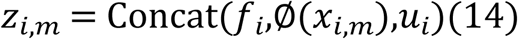

where *u_i_* is optional. When this branch is disabled, the input reduces to:

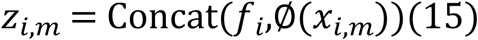

where *f_i_* denotes the multiscale fused visual representation, *u_i_* denotes the optional spatial prior feature, and ∅(*x_i_*_,*m*_) denotes the spatial encoding of the sampled coordinate.

Based on the above input, the model employs a conditional neural field *F_θ_* for expression prediction:

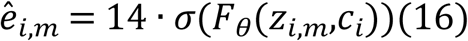

where *c_i_* denotes the conditional vector, and σ(·) denotes the sigmoid function. The constant 14 represents the unified output upper bound, which is used to constrain the predicted values to the interval [0,14]. Finally, multiple sampled predictions corresponding to the same spatial location are averaged to obtain a stable expression estimate:

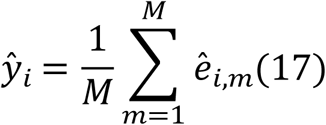

This sampling–aggregation strategy is used to improve prediction stability.

### Performance evaluation metric framework

Model performance was evaluated from three perspectives: numerical accuracy of expression abundance, spatial pattern consistency, and tissue-level topological fidelity. Predictions and ground-truth targets were clipped to [0,14] before evaluation. MSE and MAE were computed in log-transformed space, whereas PCC and HEG_PCC were computed in clipped expression space. The final prediction was obtained by averaging Monte Carlo samples (M=16).

### Expression abundance error and spatial pattern consistency

Let N denote the total number of spots in a tissue section and G denote the total number of genes. *y_ij_* and 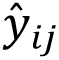 represent the ground-truth and predicted values, respectively, of the j-th gene at the i-th spot. The clipped ground-truth and predicted values are first defined as:

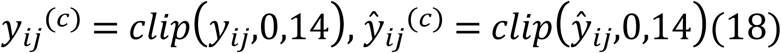

In the following text, ACG_PCC has the same meaning as PCC (ACG) in the tables, and HEG_PCC has the same meaning as PCC (HEG).

λ Mean Square Error (MSE) and Mean Absolute Error (MAE)

In this study, MSE and MAE are defined in the log(1 + *x*)-transformed space:

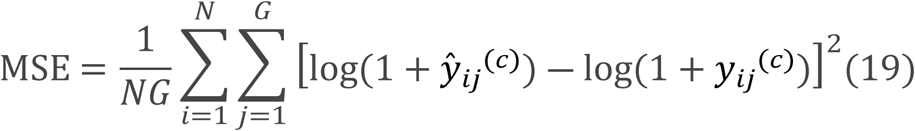

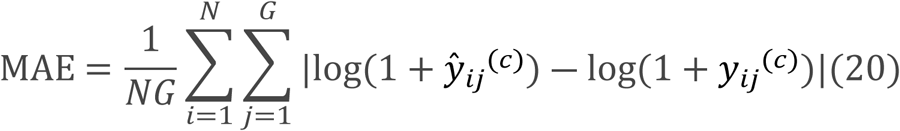

Smaller values indicate better agreement; MSE is more sensitive to large deviations, whereas MAE reflects average error.

λ Gene-wise Pearson correlation coefficient (PCC)

To evaluate the model’s ability to recover spatial expression structures, we compute the Pearson correlation coefficient for each gene in the clipped expression-value space:

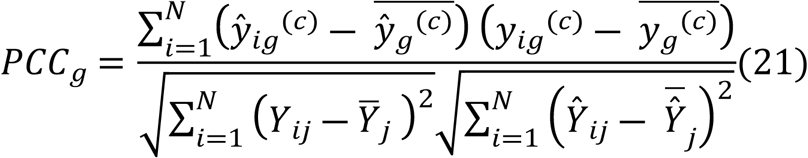

Where 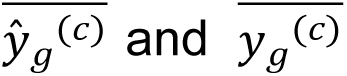 denote the mean predicted value and the mean ground-truth value of gene *g* across all spots, respectively. The overall metric is then obtained by averaging over all genes:

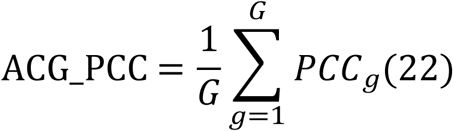

PCC reflects spatial pattern consistency rather than absolute abundance error. Its value ranges from [-1,1].

λ Highly Expressed Gene Spatial Correlation (HEG_PCC)

To evaluate the model’s ability to recover key high-signal genes, the top 50 genes with the highest mean expression levels were selected to form the set ℊ_HEG_, and their average spatial correlation was calculated as:

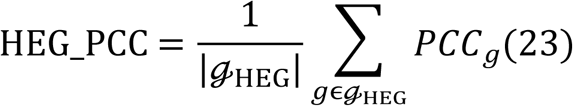

This metric evaluates recovery of spatial patterns for highly expressed genes.

### Evaluation of spatial structure and topological fidelity

Beyond the single-gene level, tissue-level reconstruction was further evaluated by applying dimensionality reduction and multiple clustering strategies to the predicted expression matrix, followed by comparison with pathologist annotations using ARI and NMI.

λ Adjusted Rand Index(ARI) ^[16]^

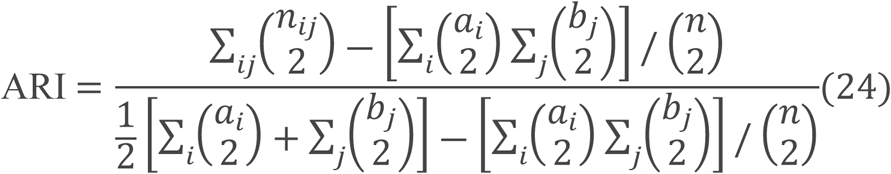

where *n_ij_* denotes the number of samples that simultaneously belong to cluster i and ground-truth class j, *a_i_* and *b_j_* denote the numbers of samples in cluster i and class j, respectively, and n denotes the total number of samples. ARI is used to measure the consistency between clustering results and ground-truth labels, with a value range of [-1,1].

λ Normalized Mutual Information (NMI) ^[17]^

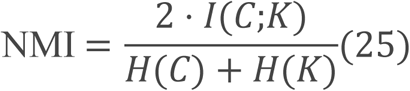

where *I*(*C*;*K*) denotes the mutual information between the clustering result C and the ground-truth annotation K, while *H*(*C*) and *H*(*K*) denote their respective entropies. NMI is used to measure the degree of shared information between two clustering results, with a value range of [0,1].

### Unified Assessment Protocol and Post-Aggregation Verification

To ensure fair comparison at the same observational scale, all methods were evaluated in spot-level observation space. For discrete baselines, spot-level predictions were compared directly with measurements. For HiFi-ST, the continuous expression field was first reconstructed and then aggregated to the spot level using the same Monte Carlo sampling mechanism as in training:

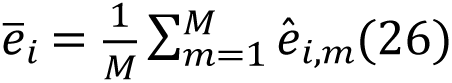

where M=16 and *σ* = 0.1. All metrics were computed from the aggregated predictions. This procedure is referred to as verify-after-aggregation.

### Baseline models

In this study, five representative baseline methods were selected:

HisToGene ^[7]^: This method uses a Vision Transformer to perform sequence modeling on histological images, captures long-range dependencies through the self-attention mechanism, and regresses gene expression.

His2ST ^[8]^: This method explicitly models spatial adjacency relationships among spots and combines visual feature extraction with graph convolution for local structural modeling.

THItoGene ^[18]^: This method integrates dynamic convolution and capsule networks to enhance the adaptability of feature representations and the ability to model hierarchical relationships.

BLEEP ^[9]^: This method constructs a shared embedding space between images and gene expression profiles through contrastive learning, enabling zero-shot expression retrieval based on image queries.

mclSTExp ^[10]^: This method jointly models visual, spatial, and expression information as a multimodal sequence and achieves modality alignment through contrastive learning.

### HiFi-ST experimental settings

HiFi-ST was implemented in PyTorch ^[19]^ and trained with Adam ^[20]^ and a cosine annealing scheduler. The initial learning rate was set to 5e-5 for HER2+ and Alex_NatGen and 2e-5 for cSCC. Models were trained for 100 epochs with batch size 256. Dropout, weight decay, and FiLM regularization were used to improve robustness. All baselines were reproduced as closely as possible to their original implementations under the same data partitioning and evaluation protocol.

### Benchmark datasets and cross-validation strategies

We evaluated HiFi-ST on three public spatial transcriptomics datasets: HER2+ ^[21]^, cSCC ^[22]^, and Alex_NatGen ^[23]^ (Supplementary Table S1). HER2+ and cSCC were generated using Legacy ST and were used to evaluate reconstruction under sparse supervision and strong structural heterogeneity, respectively. Alex_NatGen was generated using 10x Visium and was used to assess generalization under dense sampling and cross-patient distribution shift.

To reduce bias from spatial autocorrelation, all experiments adopted slice-level partitioning, such that all spots from the same section belonged exclusively to either training or test data. Five-fold cross-validation was used for HER2+ and cSCC, whereas leave-one-out cross-validation was used for Alex_NatGen. Spots outside tissue regions or annotated as artifacts were removed during preprocessing.

## Results

### Overview of the HiFi-ST model

HiFi-ST formulates spatial expression prediction as a continuous expression field learning problem and reconstructs continuous expression distributions through a conditional neural field jointly constrained by multiscale H&E images, spatial coordinates, and conditional vectors. The model further integrates FiLM-based conditional modulation with Monte Carlo local sampling and aggregation to obtain the final spot-level expression predictions (Figure 1). Based on this framework, the following results sequentially report overall performance, single-gene spatial patterns, downstream applications, and validation in special scenarios; ablation results for key components are provided in the Supplementary Materials.

**Figure 1.**
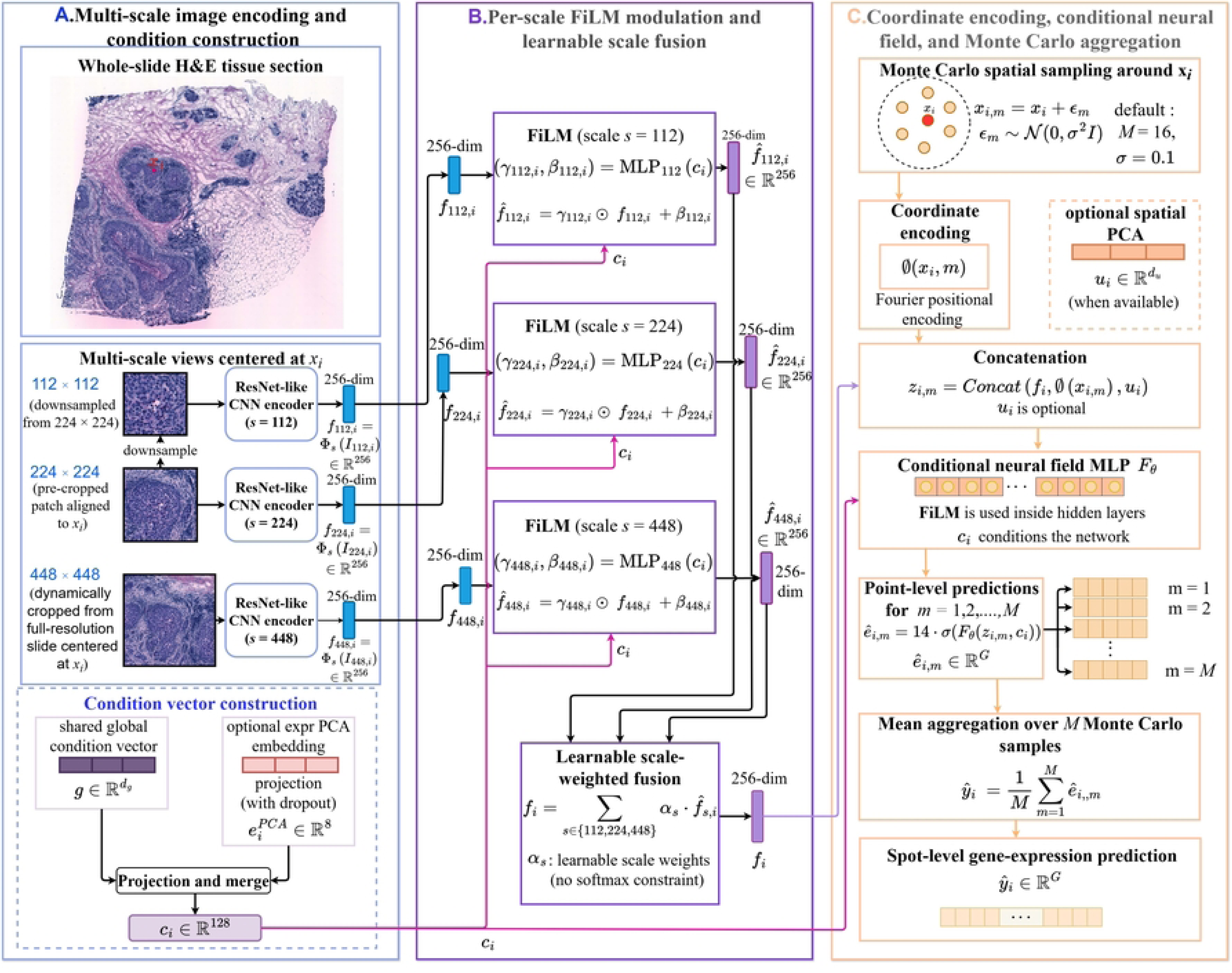
Overall framework of HiFi-ST. (A) Multiscale image encoding and conditional vector construction. (B) Multiscale conditional modulation and feature fusion. (C) Conditional neural field decoding and local sampling aggregation.

### Evaluation of High-Fidelity Continuous Field Reconstruction under Multi-level Organizational Heterogeneity

To evaluate the expression reconstruction capability of HiFi-ST under different spatial complexity scenarios, we conduct a unified comparison with HisToGene, His2ST, THItoGene, BLEEP, and mclSTExp on the HER2+, cSCC, and Alex_NatGen datasets under a slice-level data isolation protocol, as reported in Table 1. All methods are evaluated at the spot-level observation scale. For HiFi-ST, continuous field predictions are first aggregated via Monte Carlo sampling and then aligned with ground-truth spot-level expression.

**Table 1.**
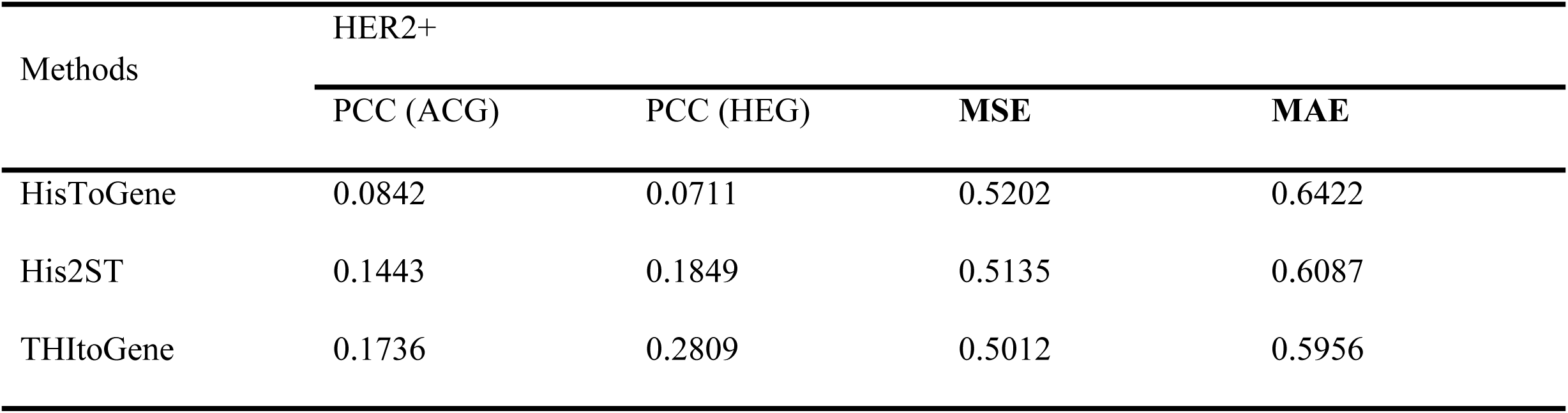
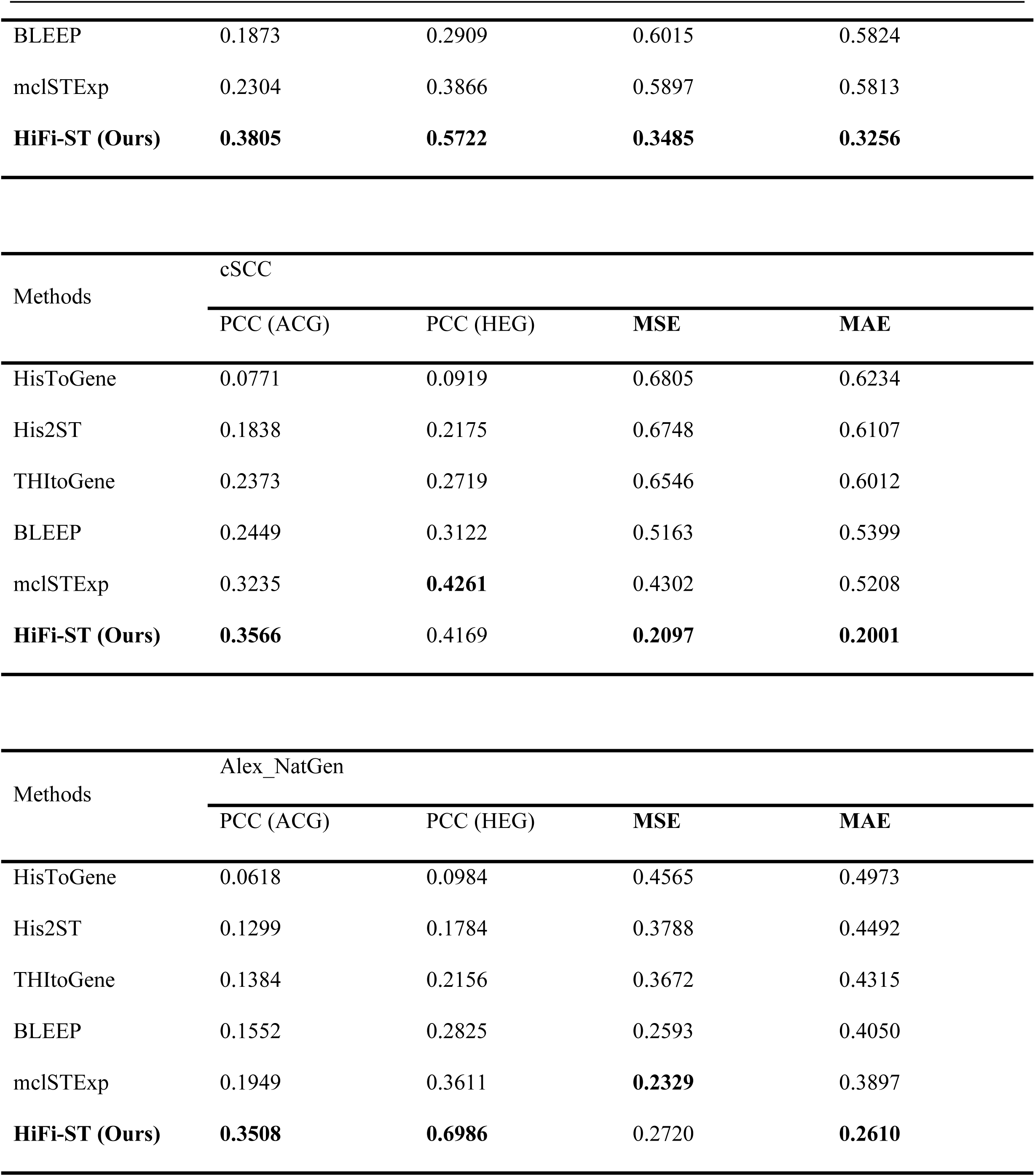
summarizes the overall performance of the various methods across the three datasets.

| Methods | HER2+ |  |  |  |
| --- | --- | --- | --- | --- |
|  | PCC (ACG) | PCC (HEG) | MSE | MAE |
| HisToGene | 0.0842 | 0.0711 | 0.5202 | 0.6422 |
| His2ST | 0.1443 | 0.1849 | 0.5135 | 0.6087 |
| THltoGene | 0.1736 | 0.2809 | 0.5012 | 0.5956 |
| BLEEP | 0.1873 | 0.2909 | 0.6015 | 0.5824 |
| mclSTExp | 0.2304 | 0.3866 | 0.5897 | 0.5813 |
| **HiFi-ST (Ours)** | **0.3805** | **0.5722** | **0.3485** | **0.3256** |

| Methods | cSCC |  |  |  |
| --- | --- | --- | --- | --- |
|  | PCC (ACG) | PCC (HEG) | MSE | MAE |
| HisToGene | 0.0771 | 0.0919 | 0.6805 | 0.6234 |
| His2ST | 0.1838 | 0.2175 | 0.6748 | 0.6107 |
| THItoGene | 0.2373 | 0.2719 | 0.6546 | 0.6012 |
| BLEEP | 0.2449 | 0.3122 | 0.5163 | 0.5399 |
| mclSTExp | 0.3235 | **0.4261** | 0.4302 | 0.5208 |
| **HiFi-ST (Ours)** | **0.3566** | 0.4169 | **0.2097** | **0.2001** |

| Methods | Alex_NatGen |  |  |  |
| --- | --- | --- | --- | --- |
|  | PCC (ACG) | PCC (HEG) | MSE | MAE |
| HisToGene | 0.0618 | 0.0984 | 0.4565 | 0.4973 |
| His2ST | 0.1299 | 0.1784 | 0.3788 | 0.4492 |
| THItoGene | 0.1384 | 0.2156 | 0.3672 | 0.4315 |
| BLEEP | 0.1552 | 0.2825 | 0.2593 | 0.4050 |
| mclSTExp | 0.1949 | 0.3611 | **0.2329** | 0.3897 |
| **HiFi-ST (Ours)** | **0.3508** | **0.6986** | 0.2720 | **0.2610** |

On HER2+, HiFi-ST achieved the best performance in both spatial correlation and error-based metrics, with ACG_PCC/HEG_PCC of 0.3805/0.5722 and MSE/MAE of 0.3485/0.3256. On cSCC, HiFi-ST achieved an ACG_PCC of 0.3566 and the best error-based metrics, while its HEG_PCC (0.4169) was close to the best-performing mclSTExp (0.4261). On Alex_NatGen, HiFi-ST showed the clearest advantage in spatial correlation, with ACG_PCC/HEG_PCC of 0.3508/0.6986; its MAE was the best, whereas its MSE was slightly higher than that of mclSTExp.

Overall, HiFi-ST showed a consistent advantage across most key metrics, particularly in spatial correlation.

### Single-gene significance and spatial pattern reconstruction

Building on the overall performance evaluation, we further examined the ability of HiFi-ST to recover fine-grained spatial expression patterns at the single-gene level. For each tissue section, gene-wise PCC was calculated across the spot dimension, and its distribution across genes was summarized (Figures 2–4). Genes containing NaN/inf values or zero variance were excluded from the statistics. Overall, HiFi-ST exhibited higher median correlations and a richer distribution of highly correlated genes across multiple datasets.

**Figure 2.**
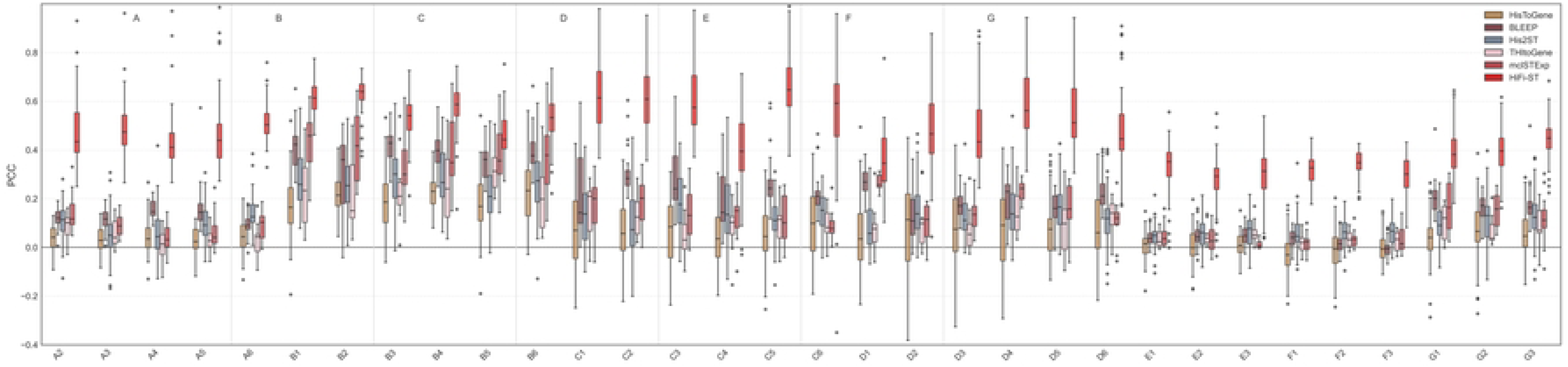
Boxplot of the gene-wise PCC distribution on the HER2+ dataset.

**Figure 3.**
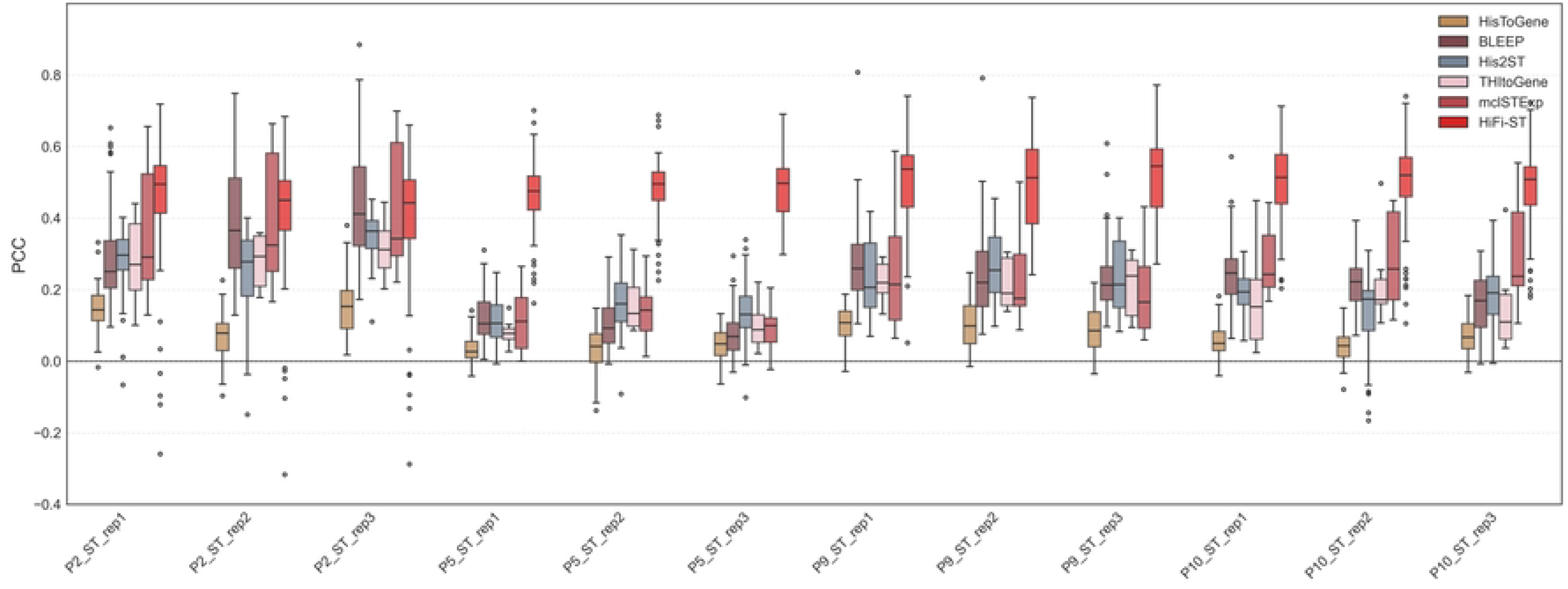
Boxplot of the gene-wise PCC distribution on the cSCC dataset.

**Figure 4.**
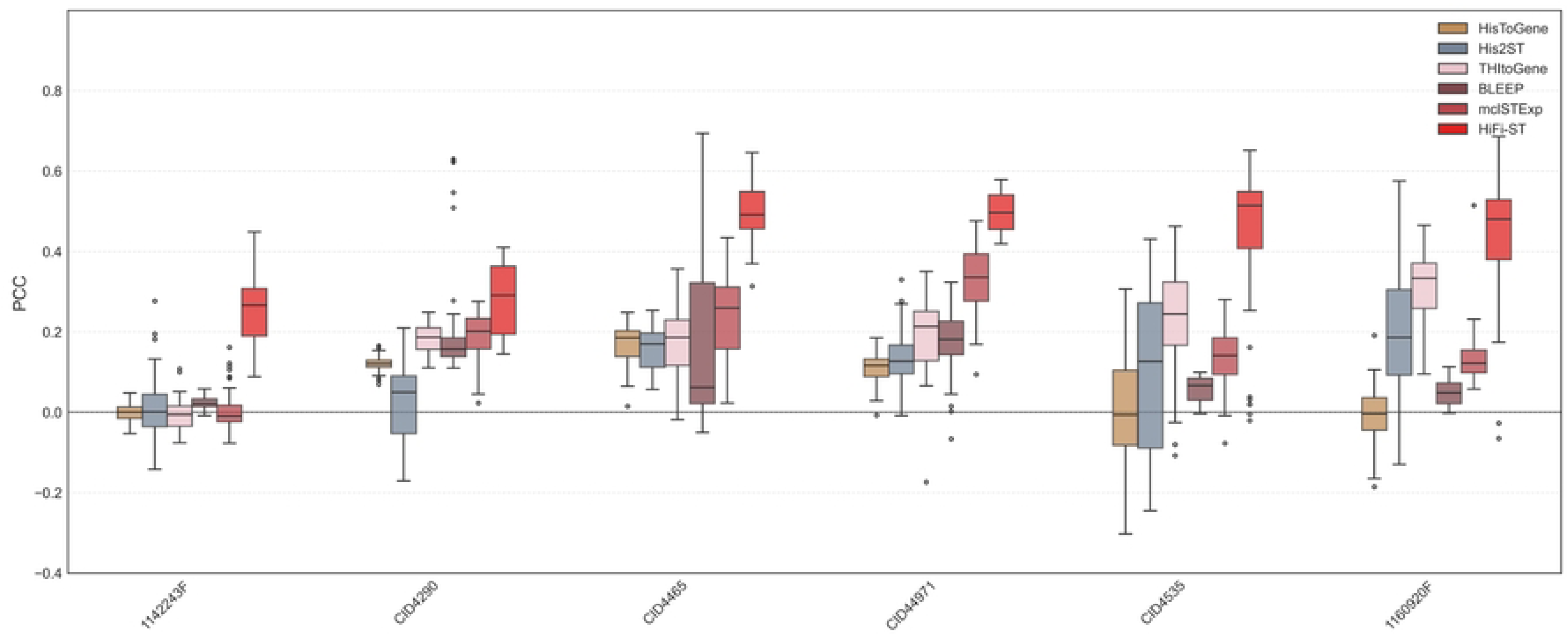
Boxplot of the gene-wise PCC distribution on the Alex_NatGen dataset.

Furthermore, for each tissue section, we performed a Pearson correlation test for each gene and defined the significance score based on the corresponding *p* value as:

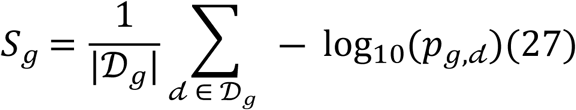

where *S_g_* denotes the set of computable tissue sections for the corresponding gene. High-confidence genes extracted based on this metric are listed in Supplementary Tables S4–S5, and Table 2 provides representative high-confidence genes identified by each method on the HER2+ dataset.

**Table 2.**
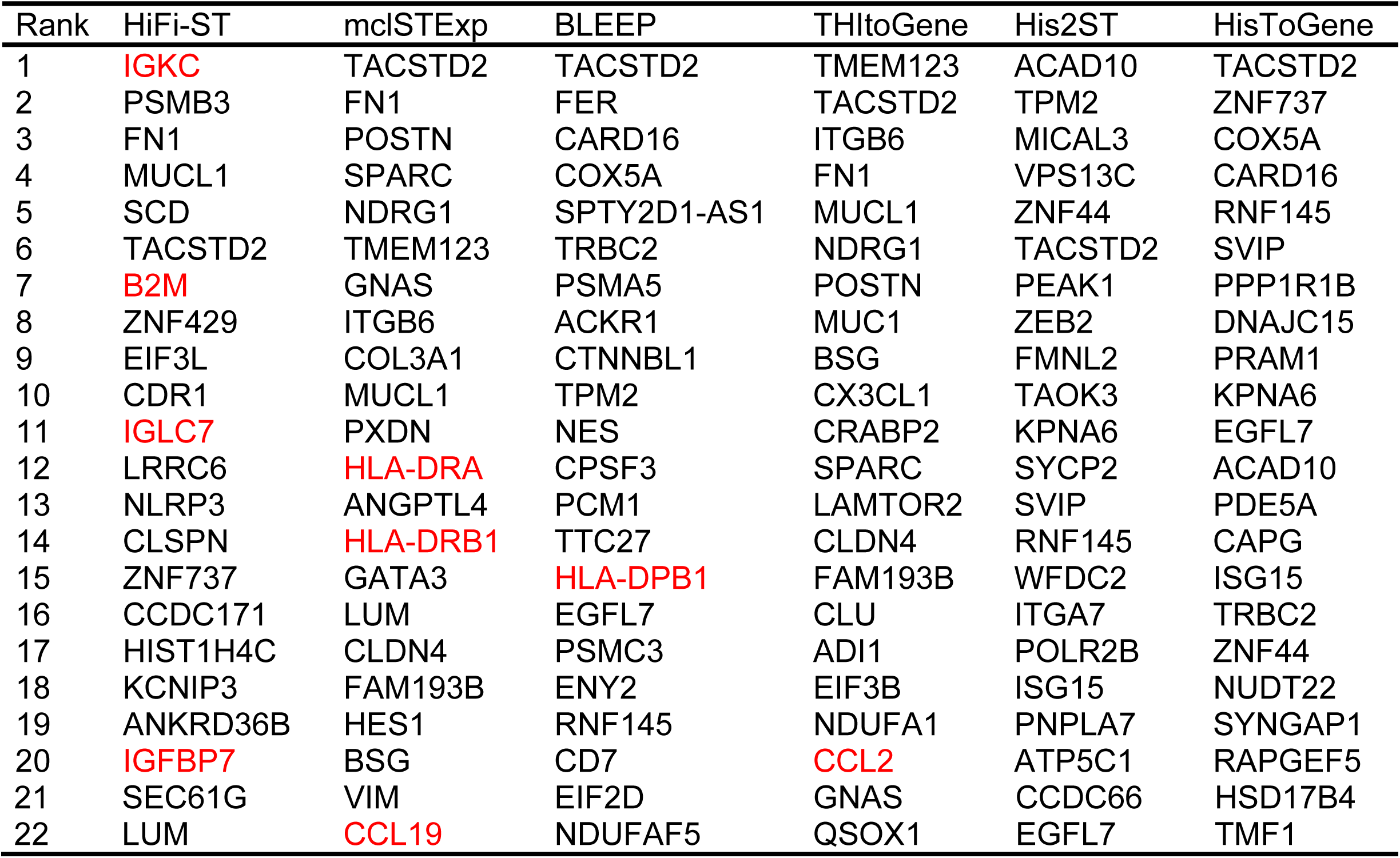
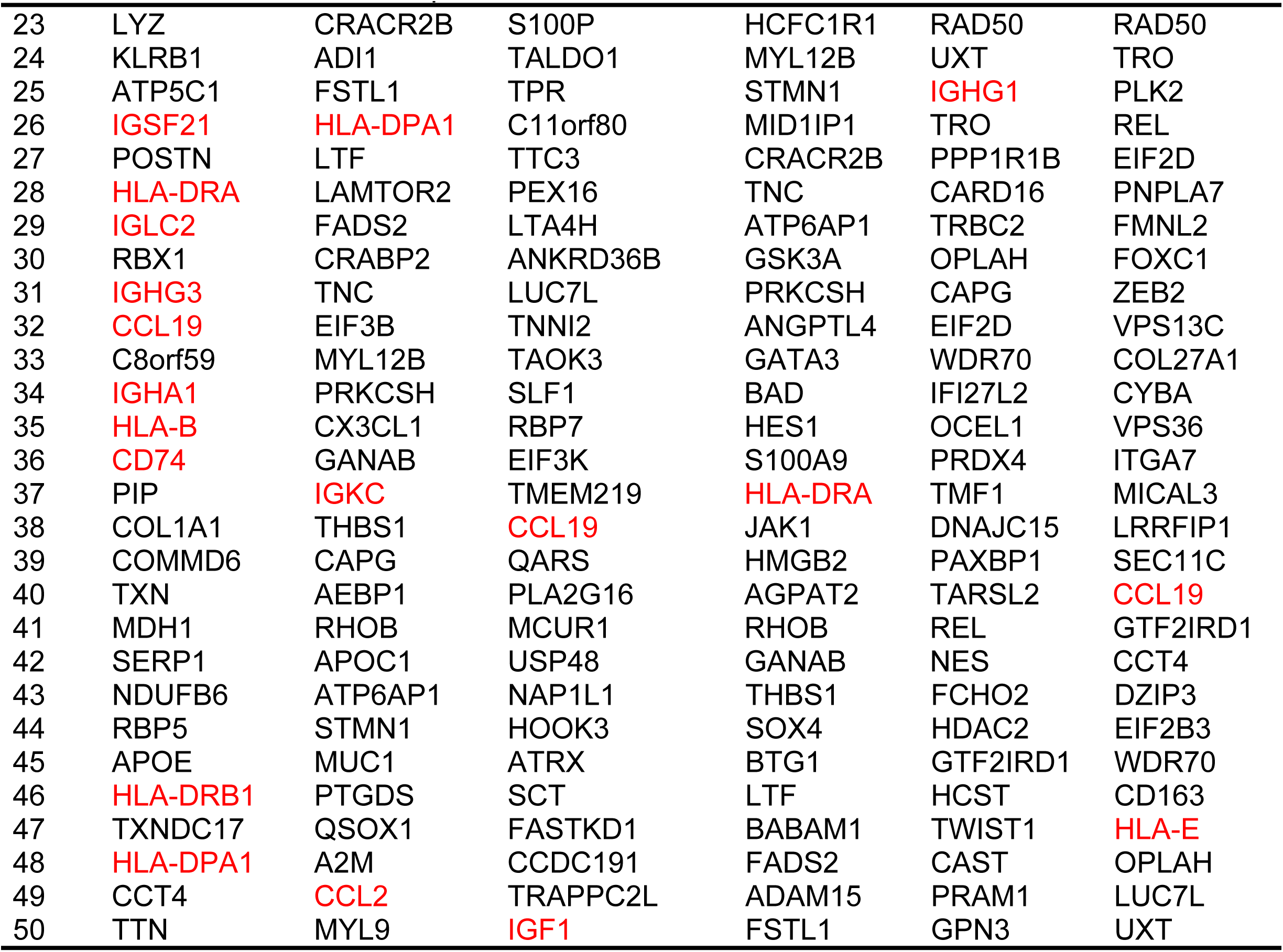
Top 50 genes ranked by prediction significance in the HER2+ dataset.

Figure 5 shows representative spatial expression visualizations. It should be noted that the PCC values shown in the figure are all slice-level PCC values for specific gene–section combinations, rather than averages across sections (Supplementary Figure S1). For genes such as FN1^[24]^, IGKC ^[25]^, and HLA-DRA ^[26]^, HiFi-ST predictions are more consistent with ground-truth spatial distributions.

**Figure 5.**
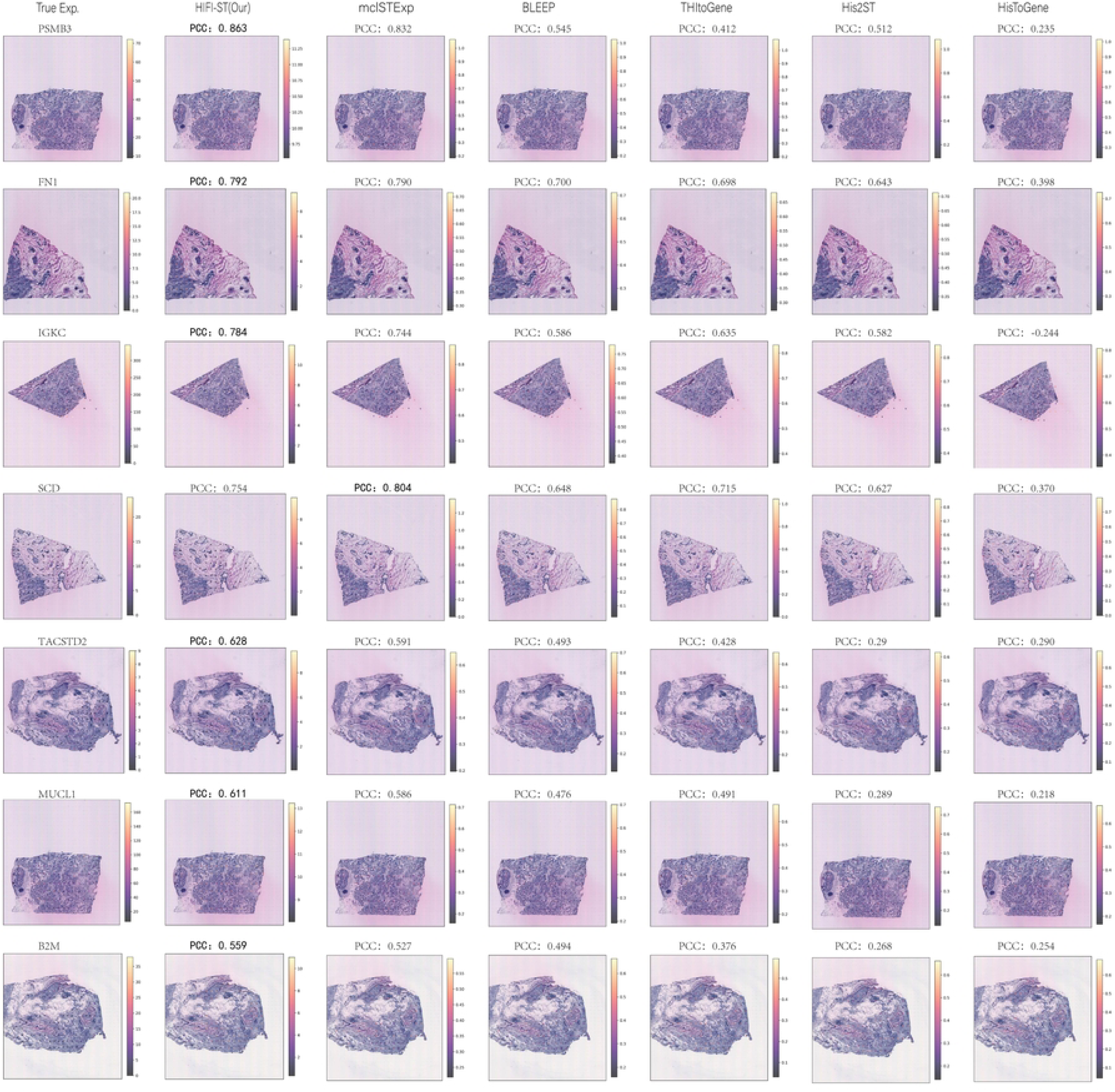
Spatial expression visualization of key genes in the HER2+ dataset.

In summary, HiFi-ST shows more stable reconstruction of key functional spatial patterns.

### Assisted identification of candidate TLS regions using multiscale representations

To assess the downstream utility of HiFi-ST representations, we evaluated candidate TLS region identification ^[27, 28]^. This was an independent within-sample supervised task and was not included in the slice-level generalization comparison.

Three image scales (112×112, 224×224, and 448×448) were encoded by the HiFi-ST multiscale encoder, followed by an MLP classification head. kNN-based relative displacement features (k=5) were used as auxiliary inputs. TLS annotations from pathologists ^[29]^ were used for supervision, with TLS_2_cat =’TLS’ as the positive class. Thresholds were selected on the validation set by maximizing F1-score. As summarized in Table 3, the detector achieved high recall but limited precision, indicating that it is more suitable for candidate-region prioritization than for replacing manual diagnosis (Figure 6).

**Figure 6.**
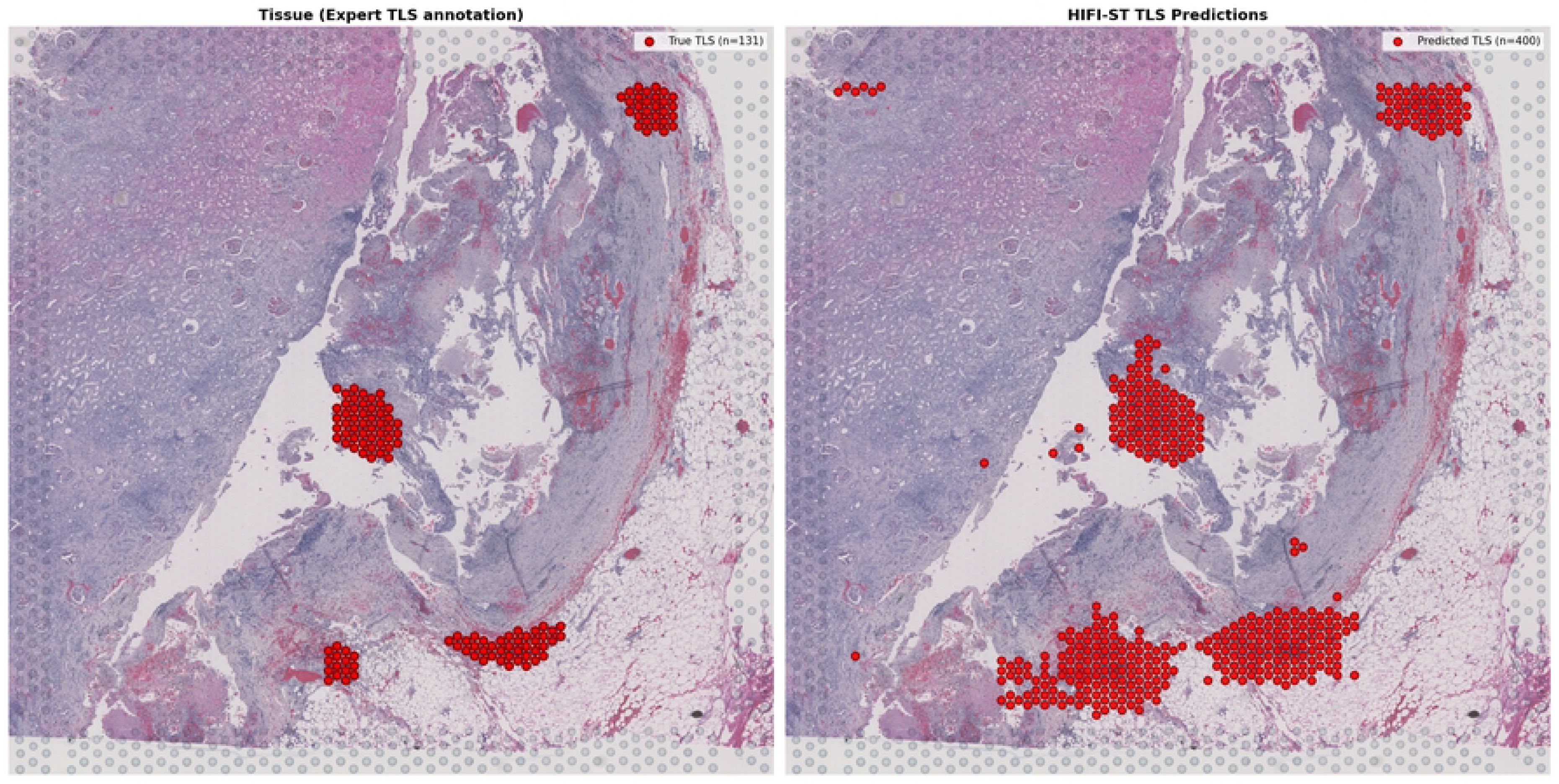
Spatial correspondence between model predictions and expert annotations.

**Table 3.** TLS candidate region identification based on HiFi-ST encoded features.

| Sample ID | Recall | Precision | F1-Score |
| --- | --- | --- | --- |
| A | 0.88 | 0.23 | 0.37 |
| B | 0.61 | 0.17 | 0.12 |
| C | 0.57 | 0.13 | 0.16 |
| D | 0.66 | 0.40 | 0.50 |
| Mean value | 0.68 | 0.23 | 0.29 |

### Fidelity-preserving molecular profile decoding under extreme morphological variation in the cSCC cohort

The cSCC cohort, characterized by dense keratinocyte arrangement, nonlinear deformation, and local tissue mixing, served as a representative robustness test. HiFi-ST still recovered key spatially structured molecular signals, with gene-wise PCCs of 0.850 and 0.838 for FTL and SPRR2A, respectively, whereas representative baselines generally ranged from 0.2 to 0.6. Biologically, FTL is associated with oxidative stress and iron metabolism reprogramming ^[30]^, whereas SPRR2A is related to keratinization ^[31]^. The stable reconstruction of these genes suggests that HiFi-ST preserves key molecular patterns even under severe morphological perturbation and can capture molecular distributions associated with typical cSCC pathological features such as keratin pearl formation and stratum corneum thickening ^[32, 33]^ (Figures 7 and 8).

**Figure 7.**
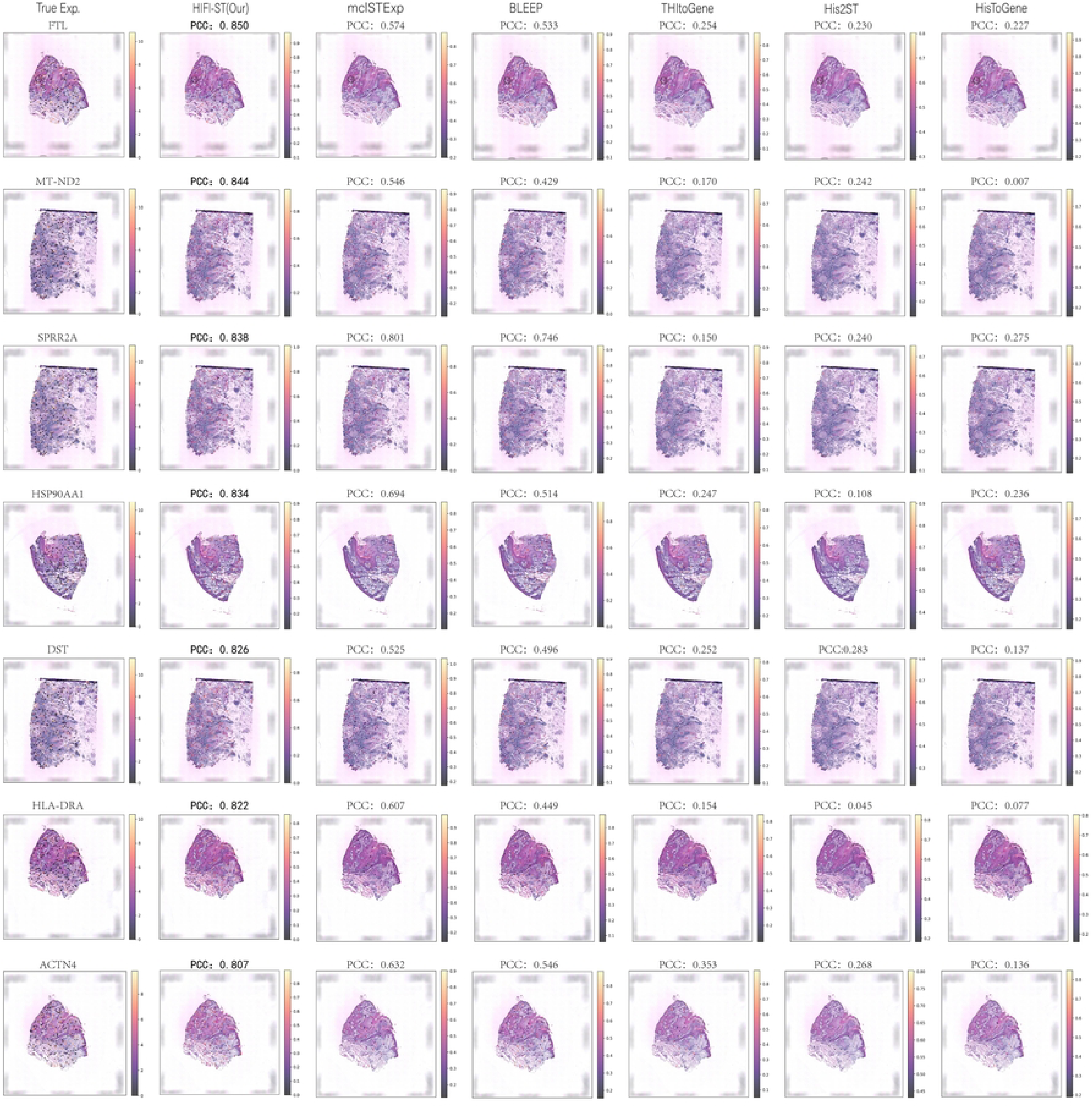
Visualization of the top-ranked genes in the cSCC dataset based on significance ranking.

**Figure 8.**
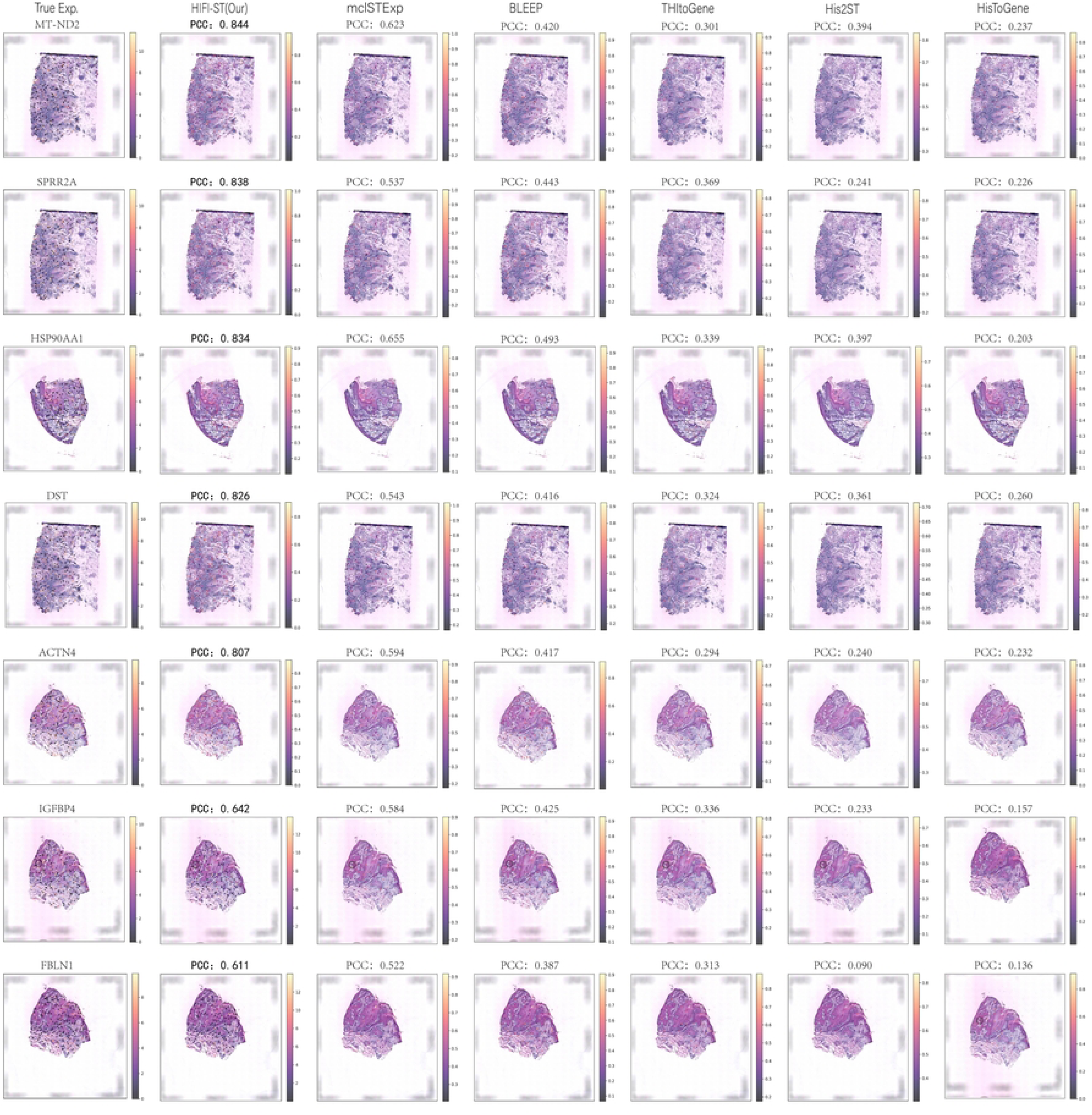
Visualization of the top-ranked genes in the cSCC dataset based on PCC ranking.

### Generalization of microenvironmental patterns across patients in the Alex_NatGen cohort

In Alex_NatGen, HiFi-ST reconstructed relatively stable spatial patterns for tumor microenvironment-related genes across patients. ECM-related genes including POSTN ^[34]^, DCN, and COL1A1 ^[35]^ remained co-localized with stroma-enriched regions across sections (Supplementary Figures S2–S3), indicating that the model captures structurally constrained spatial patterns despite cross-patient variation.

### Spatial topological domain identification based on continuous expression reconstruction

We performed unsupervised clustering on reconstructed expression matrices and evaluated consistency with reference annotations using ARI and NMI. Multiple dimensionality-reduction and clustering strategies were explored rather than a single fixed pipeline. HiFi-ST achieved higher ARI/NMI in some samples and produced more coherent boundaries in representative sections, although the benefit was not universal (Figure 9). Thus, the value of continuous reconstruction in spatial domain identification is better interpreted as a conditional gain.

**Figure 9.**
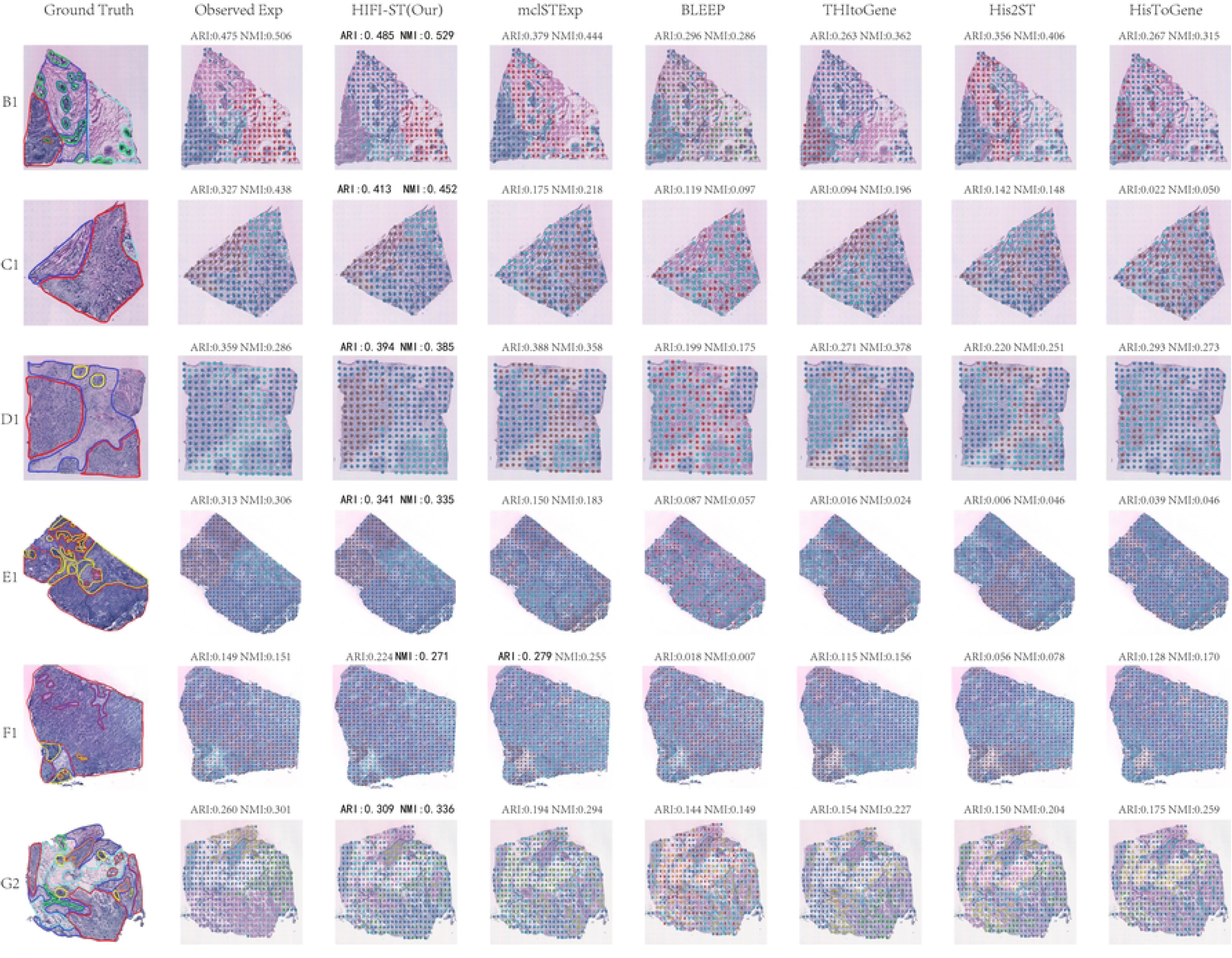
Comparison of spatial domain identification results on the HER2+ dataset.

### Ablation experiments and core component results

Systematic ablation experiments were conducted on HER2+, cSCC, and Alex_NatGen to examine the contributions of spatial encoding, multiscale modeling, conditional modulation, regional sampling and aggregation, HEG optimization, and model capacity. Overall, HiFi-ST gains arise from the synergy of high-frequency spatial encoding, multiscale conditional modulation, and regional observation modeling rather than from any single module. Detailed results are provided in Supplementary Tables S6–S13 and Supplementary Figure S5.

## Discussion

The discrete sampling mechanism of spatial transcriptomics limits the characterization of continuous expression fields. To address this issue, we propose HiFi-ST, a continuous spatial expression reconstruction framework based on conditional neural fields. Unlike conventional spot-level point regression methods, HiFi-ST integrates continuous expression field learning, spot-level regional observation modeling, and multiscale conditional modulation within a unified framework, thereby jointly accounting for expression continuity, consistency with the measurement mechanism, and cross-scale morphology–molecule mapping capability.

Experimental results demonstrate that HiFi-ST achieves overall competitive performance on the HER2+, cSCC, and Alex_NatGen datasets and outperforms baseline methods across most key metrics. In particular, the model shows consistent advantages in spatial correlation metrics (PCC and HEG_PCC), indicating improved recovery of spatial gene expression structure. Moreover, the learned continuous expression representations can support downstream spatial immunoanalysis tasks, such as assisted identification of candidate TLS regions, highlighting its potential value in tumor microenvironment analysis and spatial functional interpretation.

Despite these promising results, several limitations remain. First, the current model performs continuous modeling only in two-dimensional tissue-section space and does not explicitly incorporate three-dimensional structural information. Second, the regional integration strategy relies on uniform sampling assumptions and does not yet leverage biologically informed, non-uniform priors such as nuclei density or tissue morphology constraints. Third, TLS analysis still depends on supervised annotations, which may limit scalability in fully unsupervised settings. Finally, the study does not report uncertainty estimates stratified by tissue sections or statistical significance testing across folds, which limits the statistical rigor of cross-method comparisons.

## Conclusion

HiFi-ST provides a unified framework that elevates spatial transcriptomics from discrete spot-based observations to continuous expression field modeling. By bridging conditional neural fields with histology-guided multiscale representation learning, the model enables coherent reconstruction of spatial gene expression while maintaining consistency with the underlying measurement process.

Overall, HiFi-ST offers a principled approach for integrating morphological and molecular information in spatial omics. It demonstrates strong potential for improving spatial expression reconstruction and supporting downstream analyses such as tumor microenvironment characterization and spatial immune structure identification. This work highlights the value of continuous neural field modeling as a general paradigm for high-fidelity spatial transcriptomics inference.

## Data Availability

Details of the three public datasets used in this study are provided in the section “Benchmark datasets and cross-validation strategies.”

## Code Availability

The complete code has been made publicly available on GitHub: https://github.com/Tang-Item/HiFi-ST.git.

## Acknowledgements

The authors thank their supervisor, senior colleagues, and members of the research group for their valuable comments and constructive suggestions during the writing and revision of this manuscript.

## Author Contributions

Lei Tang designed the methodology, implemented the model, performed data processing and experiments, generated the visualizations, and drafted the manuscript. Huamei Li contributed to experimental analysis, methodological guidance, and manuscript revision. Xiaozhou Chen contributed to research conceptualization, funding acquisition, resource provision, supervision, and manuscript revision. Wenshuai Han and Xiao Yang participated in methodological discussion, experimental validation, and manuscript revision. All authors read and approved the final manuscript.

## Declarations

### Ethics approval and consent to participate

Not applicable.

### Consent for publication

Not applicable.

---

**Supplementary Figures (S1–S5) and Supplementary Tables (S1–S13) are provided in the Supplementary Materials.**

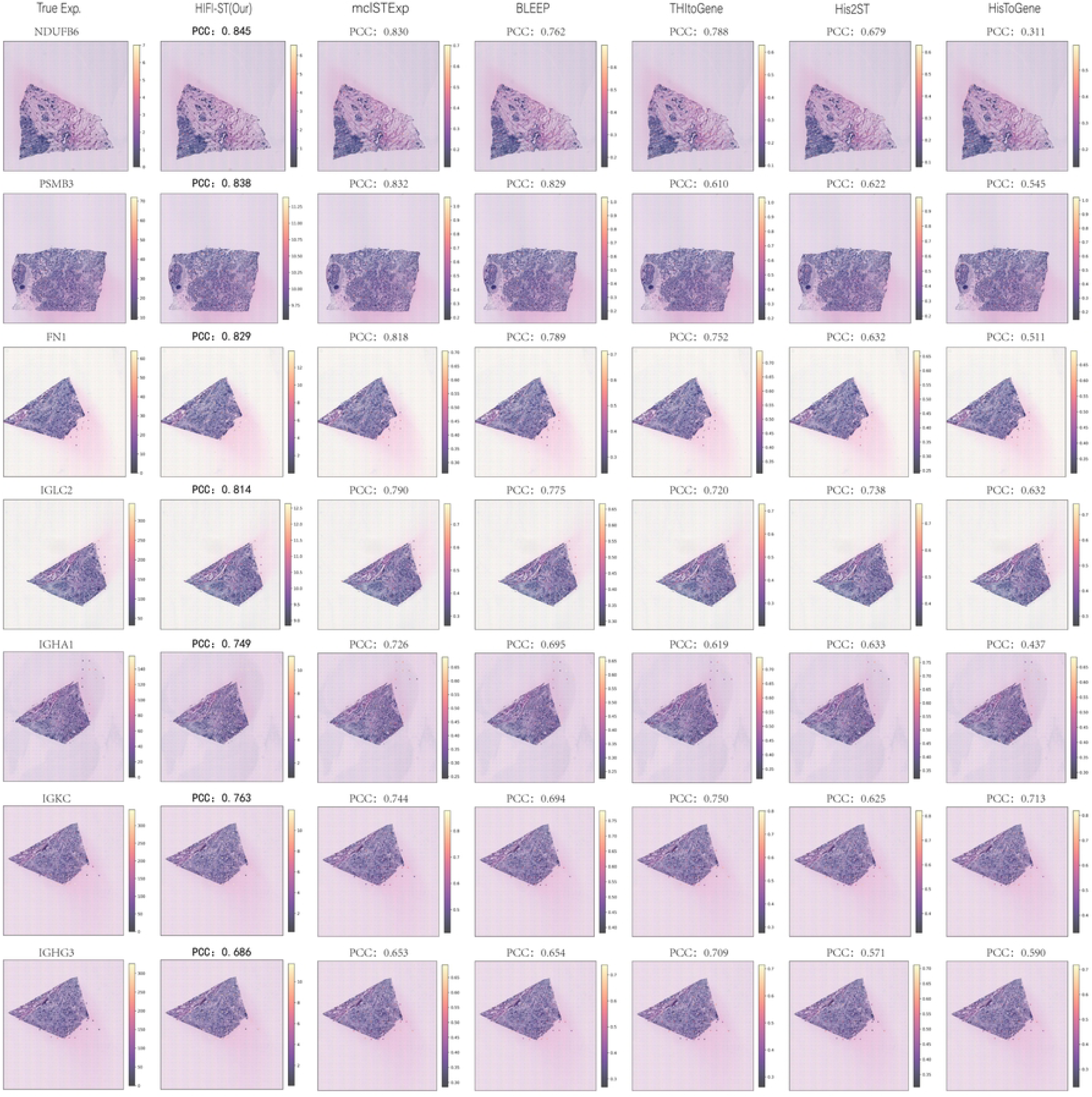

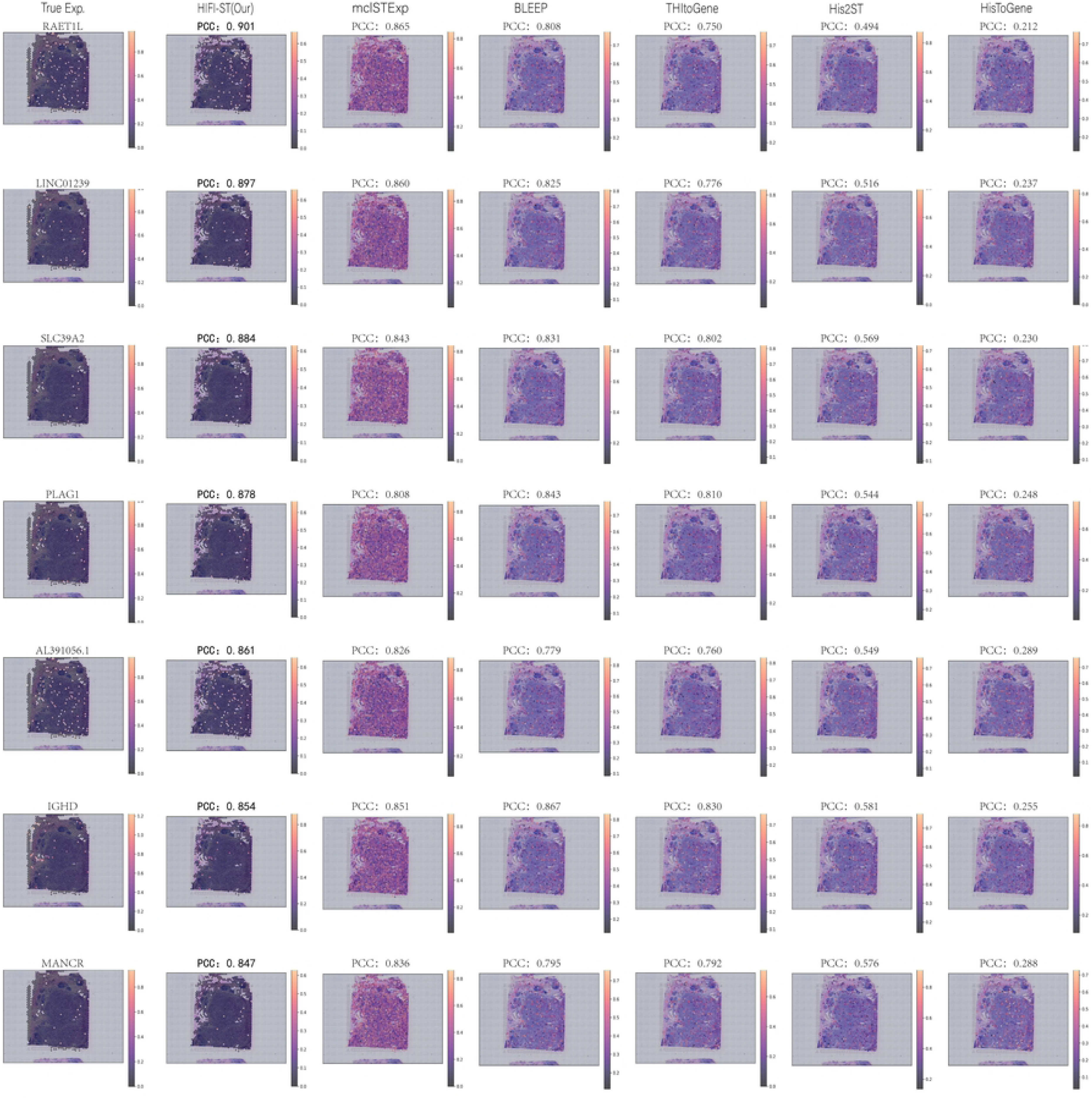

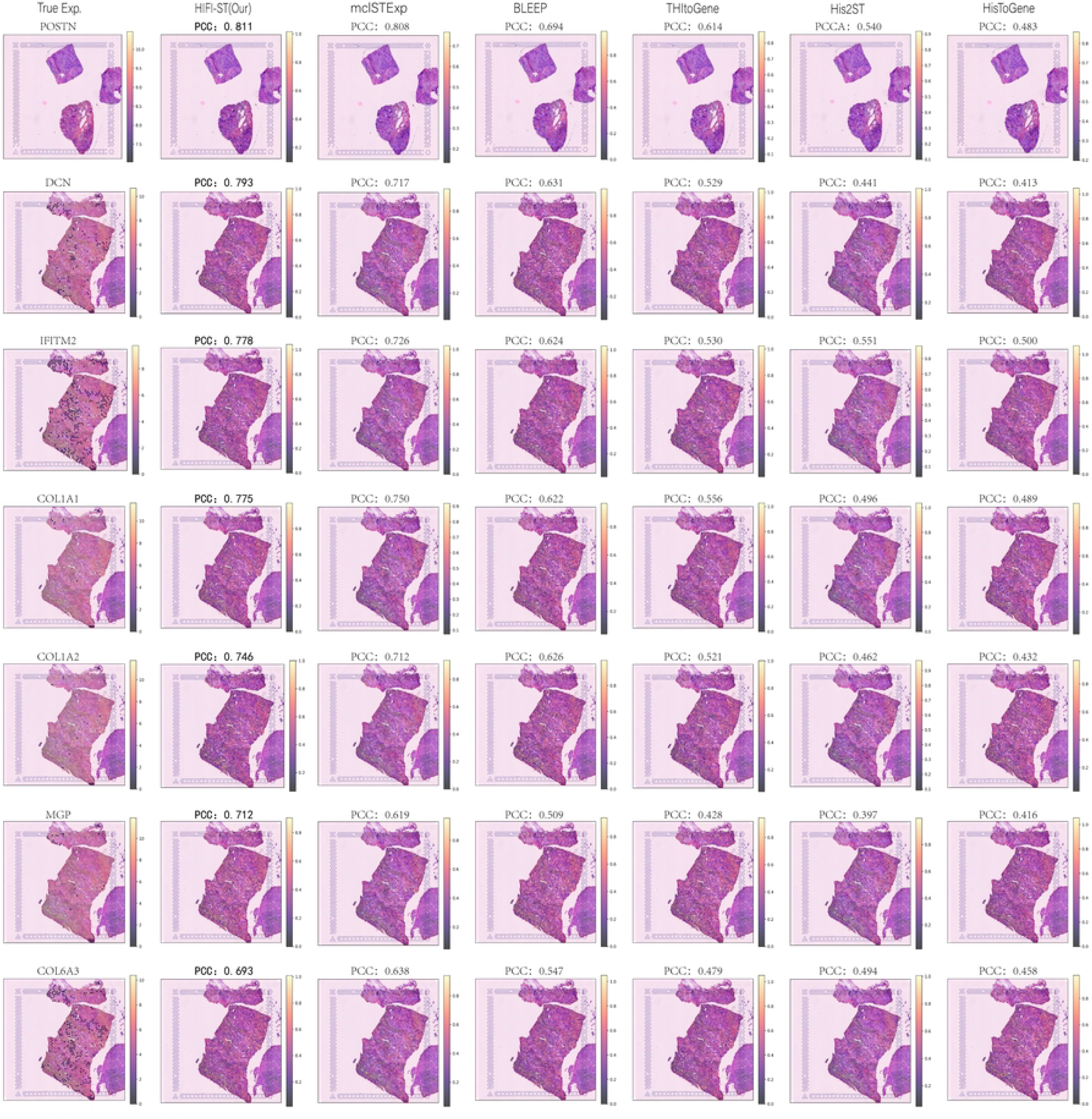

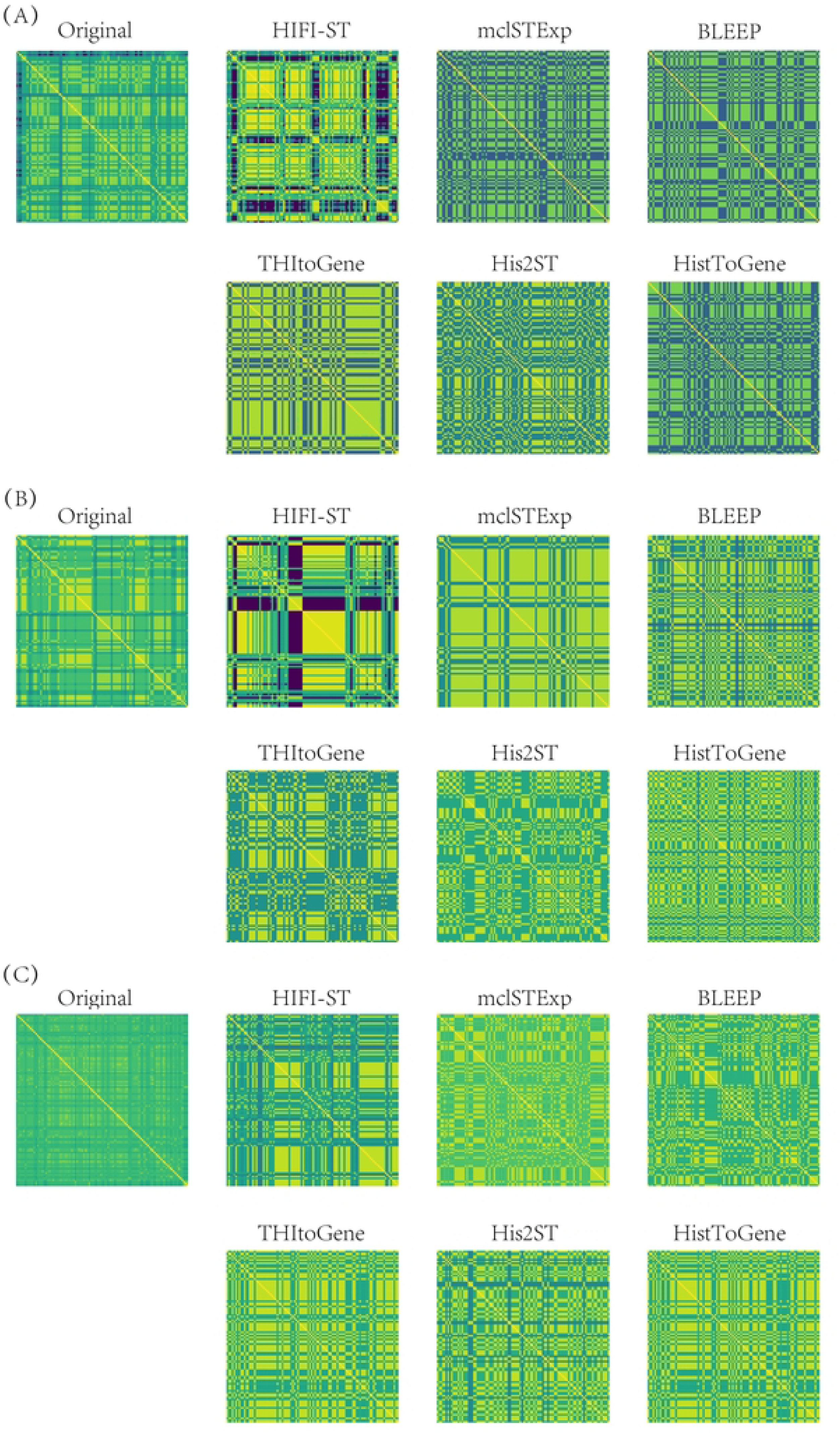

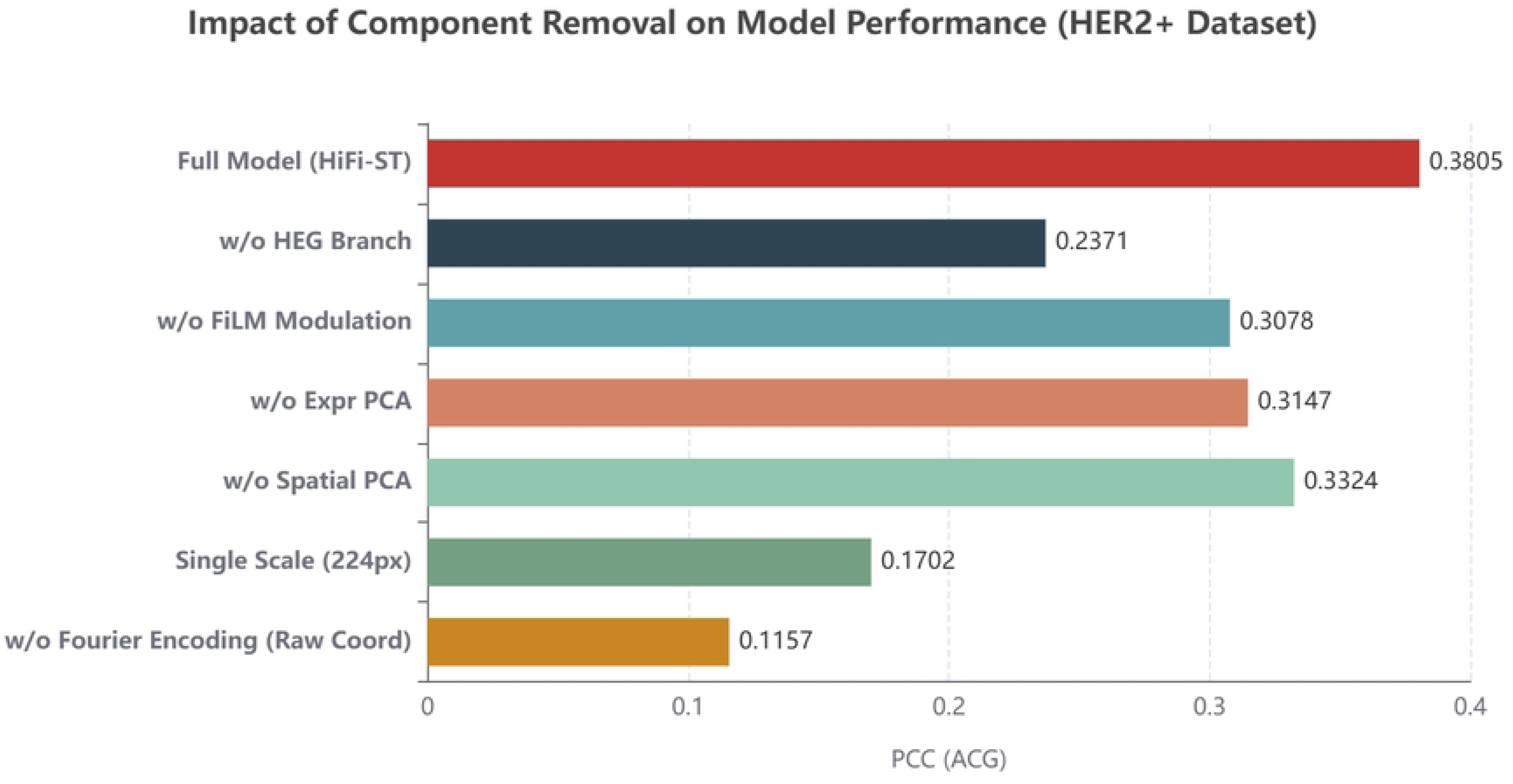

